# A *Drosophila* model for mechanistic investigation of tau protein spread

**DOI:** 10.1101/2024.04.21.590466

**Authors:** Kondalarao Bankapalli, Ruth E. Thomas, Evelyn S. Vincow, Gillian Milstein, Laura V. Fisher, Leo J. Pallanck

## Abstract

Brain protein aggregates are a hallmark of neurodegenerative disease. Previous work indicates that specific protein components of these aggregates are toxic, including tau in Alzheimer’s disease and related tauopathies. Increasing evidence also indicates that these toxic proteins traffic between cells in a prion-like fashion, thereby spreading pathology from one brain region to another. However, the mechanisms involved in trafficking are poorly understood. We therefore developed a transgenic *Drosophila* model to facilitate rapid evaluation of candidate tau trafficking modifiers. Our model uses the bipartite Q system to drive co-expression of tau and GFP in the fly eye. We find age-dependent tau spread into the brain, represented by detection of tau, but not GFP in the brain. We also found that tau trafficking was attenuated upon inhibition of the endocytic factor *dynamin* or the kinase *glycogen synthase kinase-3β* (*GSK-3β*). Further work revealed that dynamin promotes tau uptake in recipient tissues, whereas GSK-3β appears to promote tau spread via direct phosphorylation of tau. Our robust and flexible system will promote the identification of tau trafficking components involved in the pathogenesis of neurodegenerative diseases.

**SUMMARY STATEMENT:** The trafficking of toxic proteins in neurodegenerative disease is well-known but poorly understood. Our model will allow rapid and new insight into molecular mechanisms underlying this process.

## INTRODUCTION

The accumulation of brain protein aggregates is a hallmark of neurodegenerative disease (1–3). Although the composition of these aggregates, their subcellular distribution, and their precise anatomical locations vary, studies using genetic, molecular, and cell biological approaches have established that specific protein components of these aggregates are toxic and play central roles in pathogenesis (4–7). In particular, the protein tau is an essential component of the neurofibrillary tangles that mark Alzheimer’s disease, and α-synuclein is the major component of Lewy bodies in Parkinson’s disease(8, 9). Increasing evidence also suggests that neurodegenerative diseases characterized by protein aggregates share fundamental features with prion diseases (10). Prion diseases are defined by their infectious nature and the fact that prions can spread between tissues (11). While there is little evidence that other neurodegenerative diseases are infectious, an accumulating body of work indicates that the aggregates that characterize Alzheimer’s disease (AD), Parkinson’s disease (PD), Huntington’s disease, and many other neurodegenerative diseases can move between tissues and thereby promote the spread of neurodegeneration (12). Both neurofibrillary tangles and Lewy bodies appear to propagate along defined neuroanatomical pathways as neurodegeneration progresses (13–15). Autopsy of PD patients who had received fetal brain grafts revealed Lewy bodies in the graft tissue, suggesting that Lewy body pathology had propagated from the surrounding tissue (16). Furthermore, injecting aggregated forms of tau or α-synuclein into the mouse brain recruited normally folded forms of these proteins into aggregates, which were later detected in brain regions far from the sites of injection (17–20).

While recent work has begun to decipher the mechanisms by which toxic proteins spread in neurodegenerative disease, our understanding of these processes is far from complete (21). A major factor limiting progress is the lack of model systems that permit rapid analysis of candidate trafficking components. To address this matter, we have created a transgenic *Drosophila* strain that expresses tau, a toxic protein that aggregates in Alzheimer’s disease and related disorders (22). Although a number of transgenic lines have been created to study tau toxicity in *Drosophila*, these lines and the methods that have been used to monitor toxic protein trafficking have several important limitations (23–29). Our new transgenic line, coupled with a rapid and simple detection method to detect tau spread, remedies these limitations. Our transgene, expressed under the control of the Q system (30), consists of the coding sequences of human tau and GFP with an intervening T2A protein cleavage sequence (31). GFP, which is cleaved from tau during translation and does not spread, marks the expression sites of our constructs; the spread of tau can thus be detected as locations where tau is present but GFP is not.

We found that expressing our *tau-T2A-GFP* construct in the fly eye resulted in abundant tau and GFP expression in the eye as expected. Tau protein was detected in the brain in increasing abundance over time, but GFP was never detected outside the eye, indicating that tau but not GFP spread to the brain. We also found that genetic perturbations that reduce the activity of the kinase glycogen synthase kinase-3β (GSK-3β) or the endocytic factor dynamin both resulted in reduced tau trafficking. These effects were specific to tau; the same perturbations had no effect on the spread of α-synuclein, the toxic component of the Lewy body aggregates that characterize Parkinson’s disease (32). Targeted perturbations of GSK-3β and dynamin in tissue subsets further showed that GSK-3β promotes tau spread by hyperphosphorylating tau, whereas dynamin promotes tau spread by fostering the uptake of tau in recipient cells. Together, our findings indicate that our novel method for testing candidate tau spread modifiers will prove valuable in helping to decipher the mechanisms underlying the spread of tau, and possibly of other toxic proteins involved in neurodegenerative disease.

## RESULTS

### Generation and expression of a *tau-T2A-GFP* transgenic line

The goal of our work was to create a model system suitable for rapid screening of candidate factors that influence tau trafficking between tissues. Our system involves three features. First, we used the Q system (30) to drive tau expression such that we could independently use the GAL4 system (33) to perturb candidate trafficking components. The Q system works similarly to the GAL4 system in that the tissue specific expression of the exogenous transcription factor QF2 selectively drives the transcription of transgenes that contain a QF2 transcriptional response element. Importantly, QF2 does not recognize the GAL4 transcriptional response element and therefore does not interfere with the GAL4 expression system transgenes (30, 34). Second, to visualize tau spread while simultaneously marking the original site of expression, we generated a QF2-responsive construct consisting of the coding sequence of human *tau* tagged with *FLAG*, followed by a T2A protein self-cleavage sequence (31) and then by GFP (Fig. 1A). The T2A sequence releases GFP from tau, such that GFP marks the tissues where our transgene is expressed, while tissues that express tau independently of GFP represent tau spread. Third, to facilitate rapid screening, we developed a simple western blot procedure to detect tau spread.

**Figure 1.**
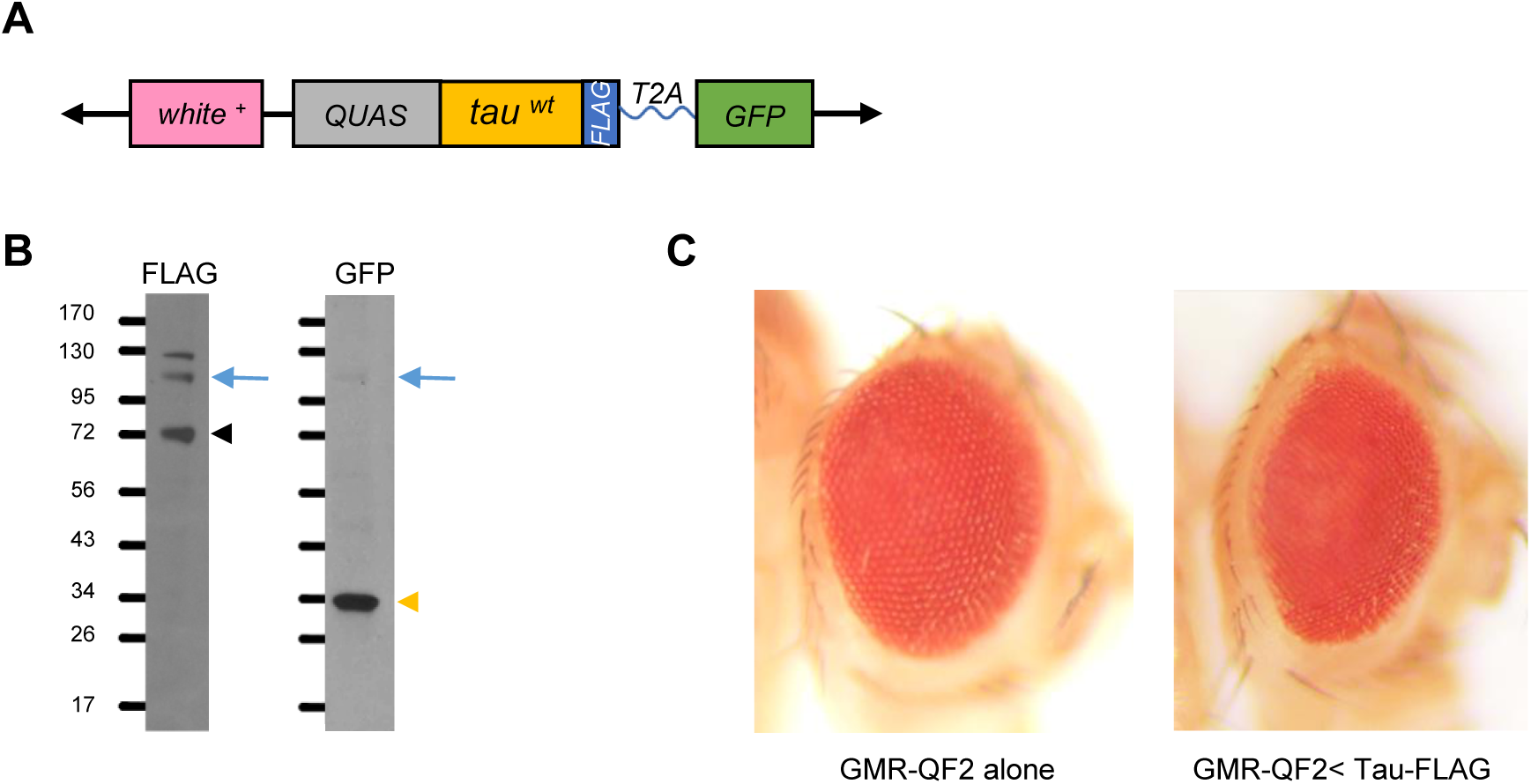
Initial characterization of the *tau-T2A-GFP* transgenic line. (A) Schematic description of the *tau-T2A-GFP* transgene. *white+* represents the marker used to detect transgenic flies. QUAS represents the Q-responsive regulatory sequence. The *tau* coding sequence also contains a C-terminal *FLAG* tag. (B) Western blot analysis of protein extracts from the heads of flies bearing the *tau-T2A-GFP* transgene and the eye-specific *GMR-QF2w* driver. Anti-FLAG antiserum (left lane) detected a tau-FLAG band of the expected size (∼70 kDa), indicated by the black arrowhead, and anti-GFP antiserum detected a GFP band of the expected size (∼30 kDa), indicated by the yellow arrowhead. Fainter bands of ∼100 kDa were also detected by both anti-FLAG and anti-GFP antisera at the expected size of the *tau-T2A-GFP* fusion protein (blue arrow), indicating that *T2A* cleavage is efficient but not complete. (C) Expression of the *tau-T2A-GFP* construct in the eye using the *GMR-QF2w* driver results in an eye that is small and mildly rough compared to the eyes of flies bearing only the driver.

After generating transgenic lines with the *tau-T2A-GFP* transgene, we crossed them to a stock bearing the eye-specific *GMR-QF2w* driver. Head protein extracts from the resulting offspring were then tested for transgene expression by western blotting for FLAG and GFP. The constructs produced appropriately sized tau-FLAG and GFP. The expected cleavage of GFP from tau appeared to be efficient, with only a minor amount of tau-GFP fusion protein detected (Fig. 1B). We also tested the effects of transgene expression using a known bioassay, disruption of the fly eye. Tau expression is toxic in many tissues and has been shown to produce a “rough eye” phenotype when expressed in the eye (23, 24). Consistent with these published findings, we found that eye-specific expression of our *tau-T2A-GFP* construct caused a mild rough eye phenotype (Fig. 1C).

### Analysis of tau spread

Previous work in *Drosophila* has shown that targeted expression of human tau protein leads to progressive spread of tau to other brain regions (26). To test whether tau expression using our constructs also led to spread beyond the site of expression, we used two approaches. First, we dissected the brains from animals expressing tau using the *GMR-QF2w* driver and performed immunocytochemistry and confocal microscopy using antisera against GFP and tau. This analysis revealed abundant tau and GFP expression in the retina as expected (Fig. 2A). Tau protein was also detected in other brain regions in the absence of GFP, indicating that tau had spread from the retina to the optic lobe (Fig. 2A). These results confirmed previously published work demonstrating that tau spreads from the fly eye to other brain regions (26). However, the methods used in previous work to detect tau spread are not suitable for rapid screening of candidate trafficking modifiers. In particular, it is difficult to quantify spread accurately using immunocytochemistry, and the process of brain dissection, immunocytochemical staining, and confocal microscopy is laborious and time-consuming. We therefore developed a novel method for detecting tau spread that was simple and rapid. We separated fly eyes from heads using a razor blade (Fig. 2B) and created separate extracts from the dissected eyes and the heads without eyes (henceforth “central heads”). Immunoblotting of these extracts revealed abundant tau and GFP in eyes, as expected for the original site of expression of the constructs (Fig. 2C). Central head extracts, on the other hand, contained tau protein but never GFP. This finding confirmed that our method was capable of detecting the spread of tau protein beyond the area in which it was expressed. The abundance of tau detected in central heads increased as flies aged, mimicking the age-dependent spread of toxic proteins in neurodegenerative disease (Fig. 2C, D).

**Figure 2.**
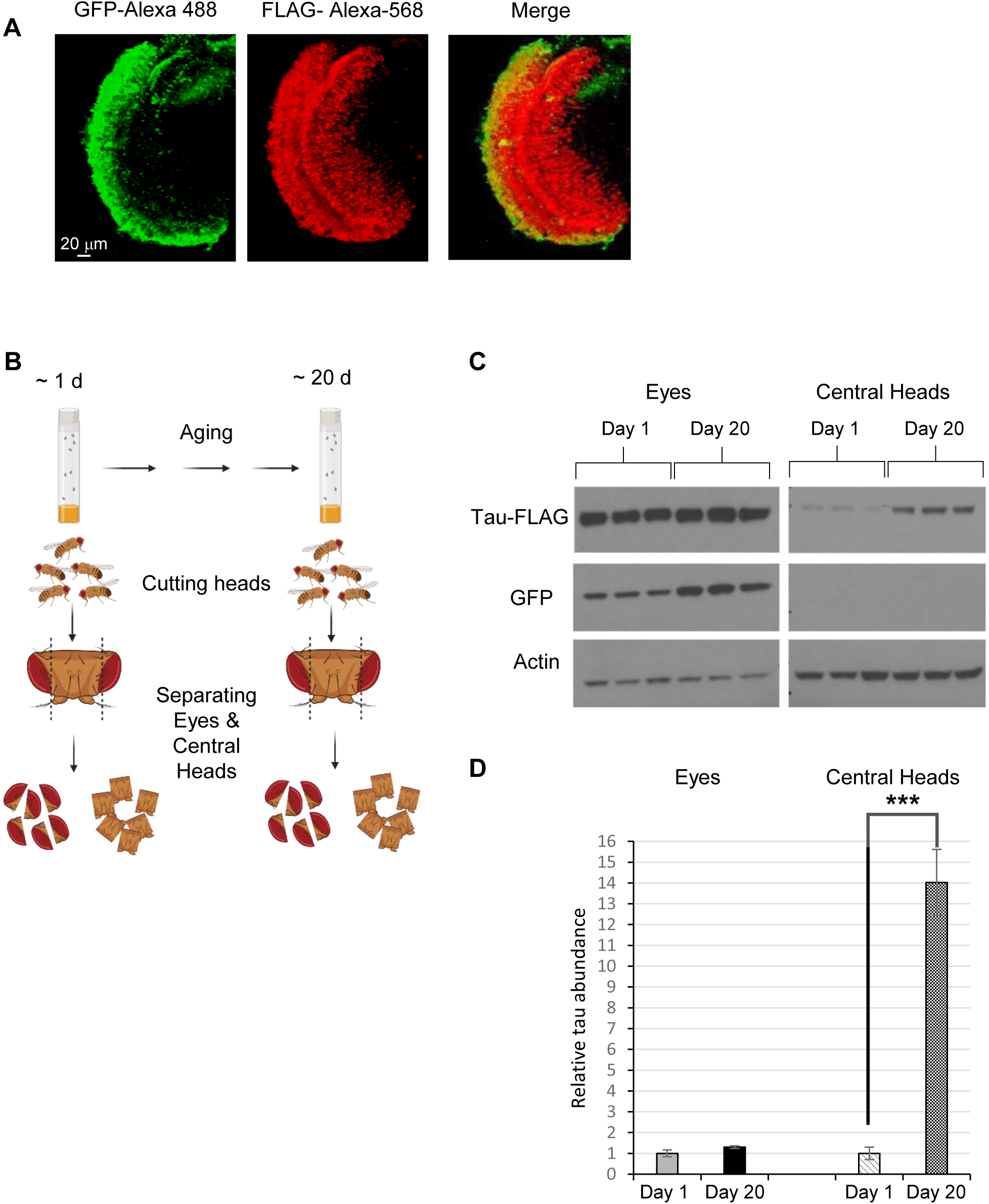
Human tau expressed in the fly eye is detected in the brain and accumulates with age. (A) Confocal images of eye and optic lobe from the brains of 40-day-old flies expressing the *tau-T2A-GFP* construct under control of the eye-specific *GMR-QF2w* driver. (B) Schematic of the experimental approach to measuring tau spread. (C) Western blot analysis using antisera against FLAG, GFP, and actin, performed on protein extracts from the eyes and central heads of 1-day-old and 20-day-old flies expressing tau in the eye. As in panel A, the flies bore both the *GMR-QF2w* driver and the *tau-T2A-GFP* construct. Each condition was represented by three biological replicates. (D) Quantification of tau abundance normalized to actin levels using the data shown in panel C. \*\*\**p* < 0.005 by Student *t-*test.

### Exploring the influence of candidate modifiers of tau spread

Having validated our system, we tested whether it could be used to identify modifiers of tau spread. One of the candidate factors we tested was *glycogen synthase kinase 3β* (*GSK-3β*). Previous work has established that hyperphosphorylation of tau contributes to its toxicity (35) and that GSK-3β is among the kinases responsible for this hyperphosphorylation (36). GSK-3β is also a modifier of tau toxicity in *Drosophila* and a modifier of Parkinson’s disease risk (23, 37, 38). Most importantly, recent work with another *Drosophila* model of tau spread has shown that inactivating GSK-3β decreases tau spread as measured by immunohistochemistry (26). To independently validate the role of GSK-3β in tau spread, we performed knockdown of this factor in flies expressing our *tau-T2A-GFP* transgene. Specifically, we used the pan-neuronal driver *nSyb-Gal4* to express RNAi targeting *GSK-3β*. We found that inactivating *GSK-3β* significantly reduced tau spread from the eye to the brain (Fig. 3A, B). Importantly, tau abundance in the eye was not affected, indicating that the reduced abundance of tau in central heads resulted from reduced spread rather than reduced expression. These results support previous findings that *GSK-3β* influences tau spread, and they establish the feasibility of our simple and rapid approach for testing candidate trafficking factors.

**Figure 3.**
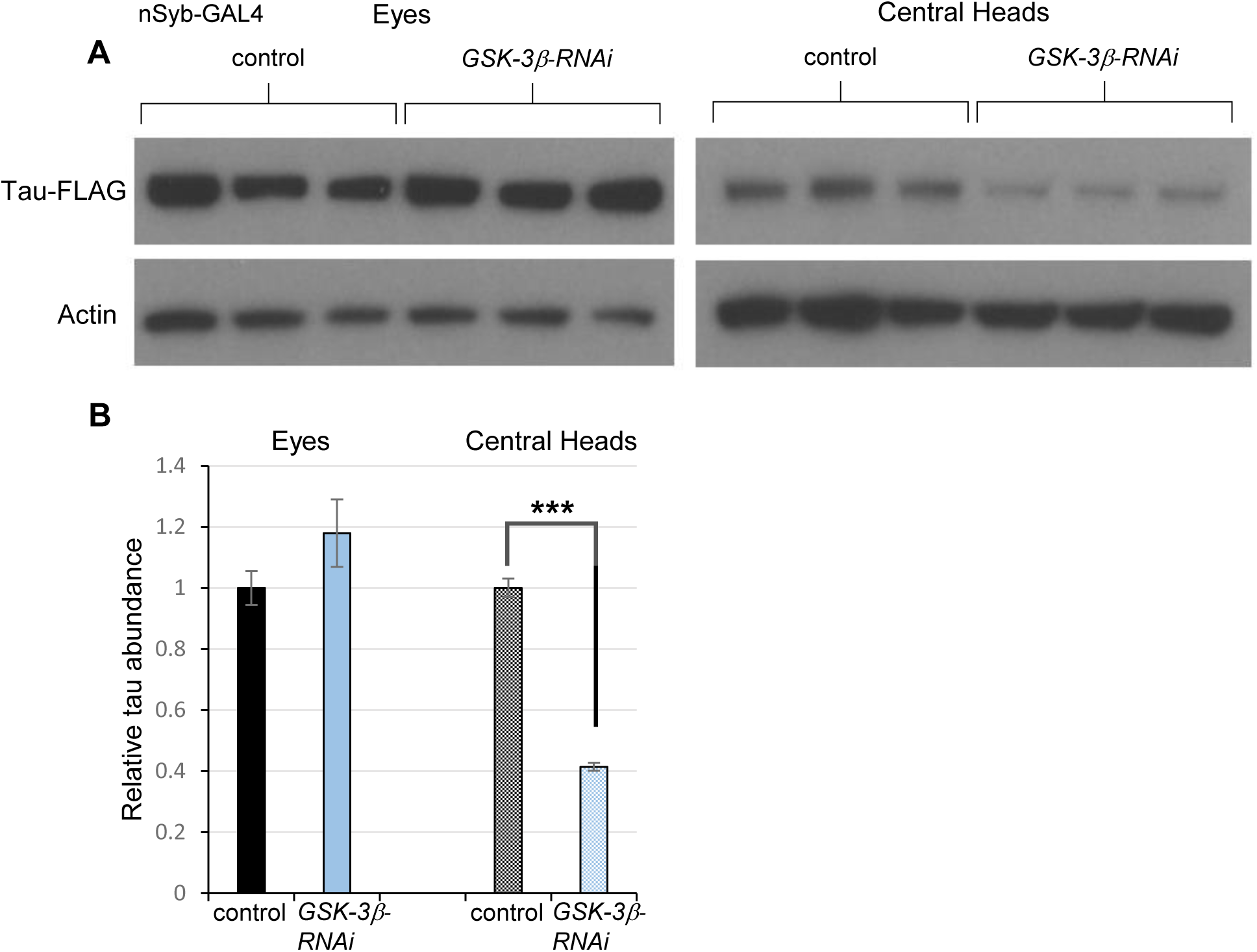
Inactivation of *GSK-3β* decreases tau spread. (A) Western blot analysis of protein extracts from the eyes and central heads of 20-day-old flies expressing tau in the eye and RNAi to *GSK-3β* in neurons. Specifically, *GMR-QF2w* drove *tau-T2A-GFP*, and the pan-neuronal driver *nSyb-GAL4* drove a GAL4-responsive RNAi targeting *GSK-3β*. Control flies lacked the RNAi transgene. Each condition was represented by three biological replicates. (B) Quantification of the data in panel A. \*\*\**p* < 0.005 by Student *t-*test.

The next candidate factor we chose to interrogate was the endocytosis factor dynamin. Work in both vertebrate and invertebrate model systems strongly suggests that endocytosis promotes the spread of toxic proteins involved in neurodegeneration by facilitating uptake of the protein in recipient cells (21, 28). We therefore tested whether inactivating dynamin in flies co-expressing the *tau-T2A-GFP* transgene in the eye would reduce the spread of tau. To perform this experiment, we used *nSyb-GAL4* to express a dominant-negative form of dynamin (39) in neurons. We found that expression of the dominant-negative dynamin transgene significantly reduced tau spread from the eye to the brain. As was the case with *GSK-3β* inactivation, tau abundance in the eye was not detectably affected by expression of dominant-negative dynamin, again indicating that the reduction of tau spread to the brain was not a secondary consequence of reduced tau expression (Fig. 4A, B).

**Figure 4.**
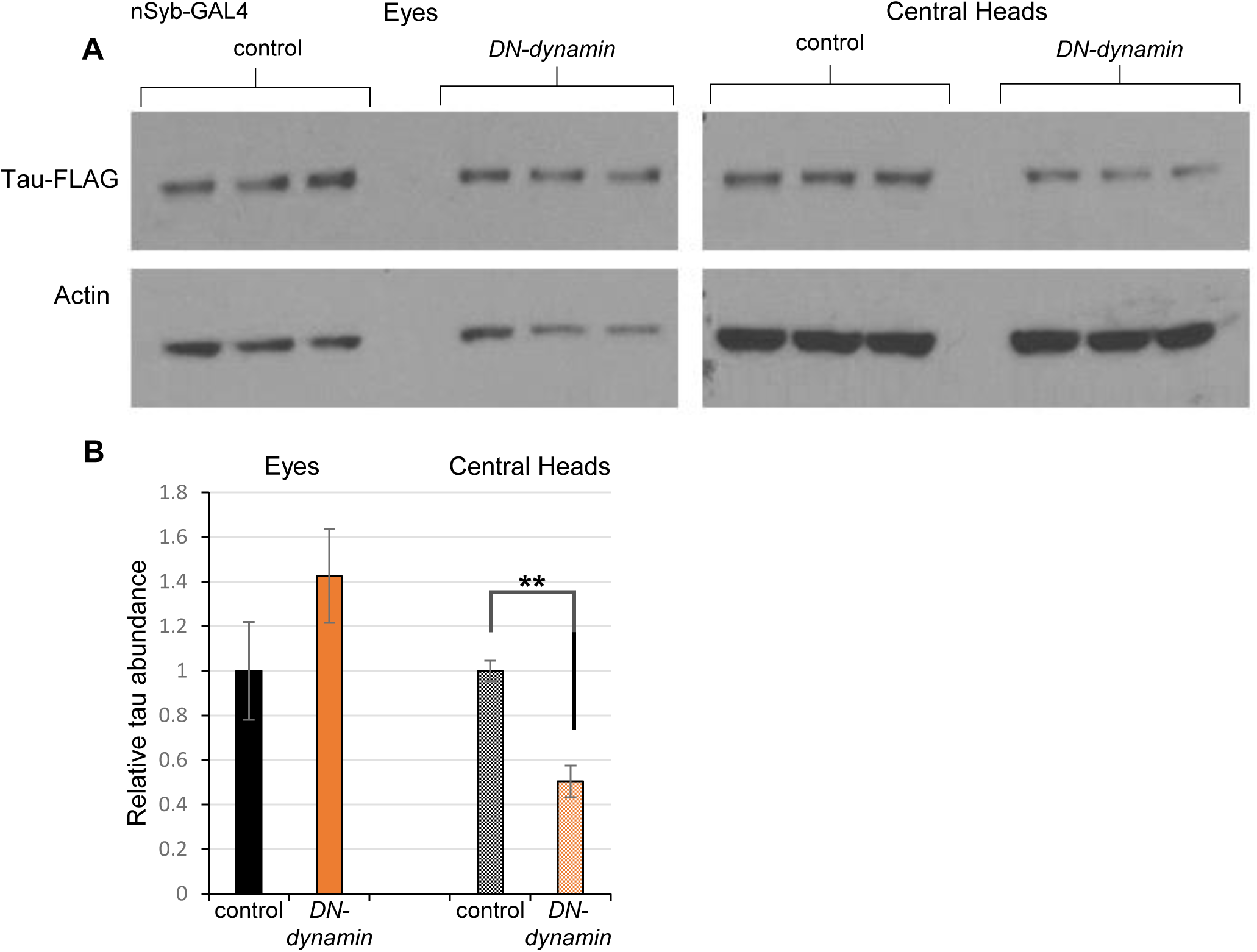
Genetic inhibition of dynamin decreases tau spread. (A) Western blot analysis of protein extracts from the eyes and central heads of 15-day-old flies expressing tau in the eye and a dominant-negative (DN) form of dynamin in neurons. The experimental group bore *GMR-QF2w* driving *tau-T2A-GFP* and *nSyb-GAL4* driving a dominant-negative dynamin. Control flies lacked the GAL4-responsive transgene. Each condition was represented by three biological replicates. (B) Quantification of the data in panel A. \*\**p* < 0.01 by Student *t-*test.

Work in model systems has suggested that tau protein spread may occur at neuronal synapses (40). Moreover, a *Drosophila* study previously showed that inactivating the *N-ethylmaleimide sensitive factor, vesicle fusing ATPase* gene (*NSF*), which plays a critical role in synaptic vesicle recycling, reduced the spread of huntingtin protein (28). Thus, we tested whether using the pan-neuronal *nSyb-Gal4* driver to express RNAi against *Drosophila* NSF would reduce the spread of human tau from the fly eye. The amount of tau detected in central heads was greatly reduced upon inactivation of NSF (Fig. 5A, B). However, the overall abundance of tau in the eye was also severely reduced by this manipulation (Fig. 5A, B), and the extent of tau reduction was similar in the two different locations. Together, these findings suggest that the reduced spread of tau seen upon NSF inactivation is likely a secondary consequence of reduced tau expression rather than altered spread. While this finding was unexpected, it reveals a useful feature of our system: It facilitates the distinction between genetic factors that influence tau spread and those that merely appear to affect tau spread as a consequence of their effect on tau abundance.

**Figure 5.**
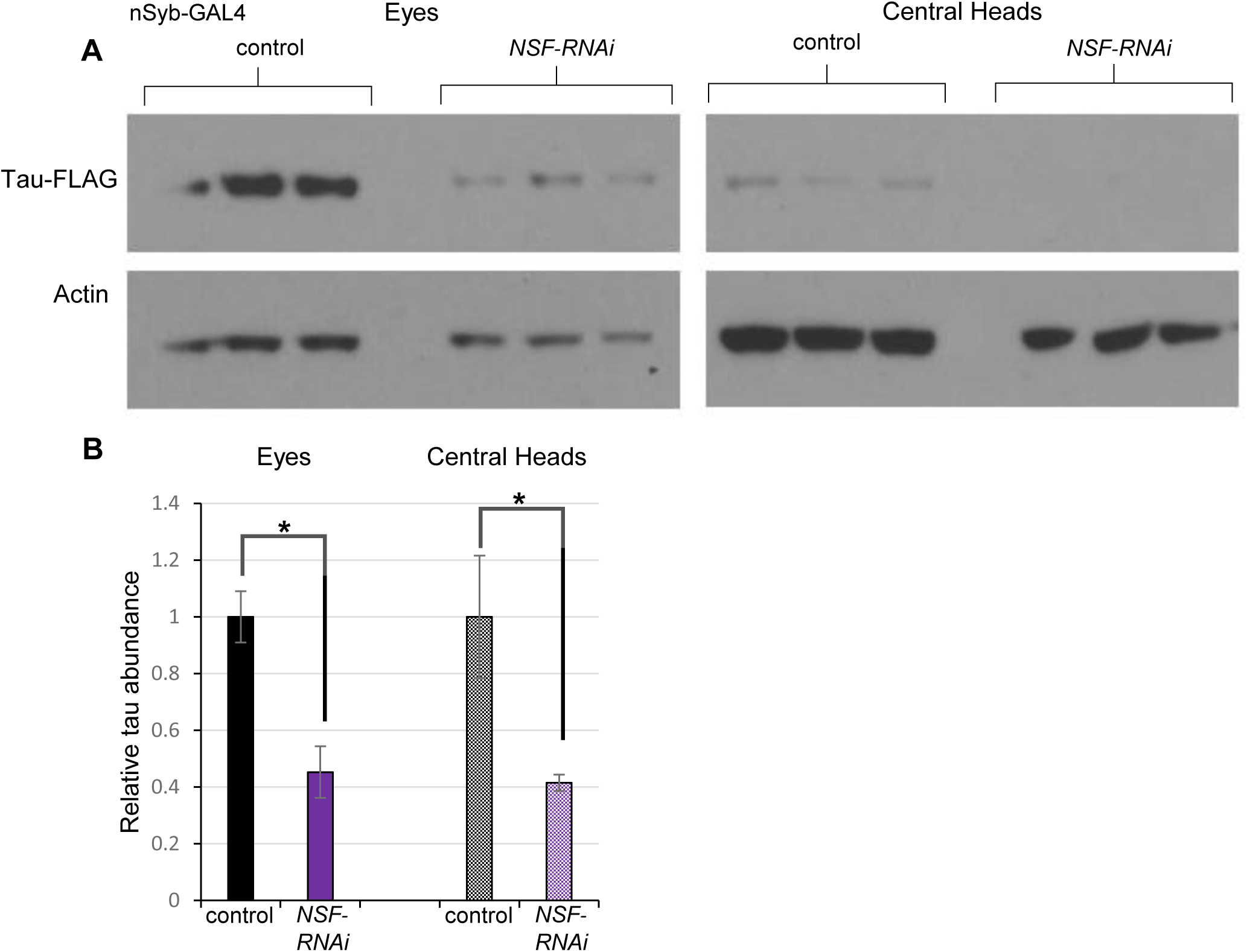
Tau abundance and spread are equally reduced upon inactivation of NSF. (A) Western blot analysis of protein extracts from the eyes and central heads of 30-day-old flies expressing *tau-T2A-GFP* in the eye and RNAi targeting *NSF* in neurons. The experimental group bore *GMR-QF2w* driving *tau-T2A-GFP* and *nSyb-GAL4* driving a GAL4-responsive RNAi to *NSF*. Control flies lacked the RNAi transgene. Each condition was represented by three biological replicates. (B) Quantification of the data in panel A. \**p* < 0.05 by Student *t-*test.

### Testing whether tau spread modifiers act cell autonomously or non-autonomously

We next sought to determine whether factors influencing tau spread acted in the cells or tissues where tau was originally expressed (source tissues), or in the cells or tissues to which tau spread (recipient tissues). To distinguish between these possibilities, we repeated our genetic perturbations, this time using the *GMR-GAL4* driver to target our perturbations to the eye. The dominant-negative dynamin construct, which reduced tau spread when driven in neurons, had no effect on tau spread when driven in the eye (Fig. 6A, B). However, eye-specific expression of the *GSK-3β* RNAi caused a decrease in tau spread (Fig. 6C, D) that was comparable to the decrease seen with *nSyb-GAL4* (Fig. 3). To further explore the influence of *GSK-3β* on tau spread, we used the *GMR-GAL4* driver to express a GAL4-responsive *GSK-3β* transgene in the eyes of tau-expressing flies. This manipulation resulted in a dramatic increase in tau spread (>5-fold relative to controls; Fig. 6E, F). Overexpressing GSK-3β also caused a mobility shift in tau on western blot that was consistent with phosphorylation, in accordance with previous work showing that tau is a direct substrate of GSK-3β (36). Together, these findings indicate that the endocytic factor dynamin promotes tau uptake in recipient tissues, whereas GSK-3β activity promotes tau spread by hyperphosphorylating tau in source tissues.

**Figure 6.**
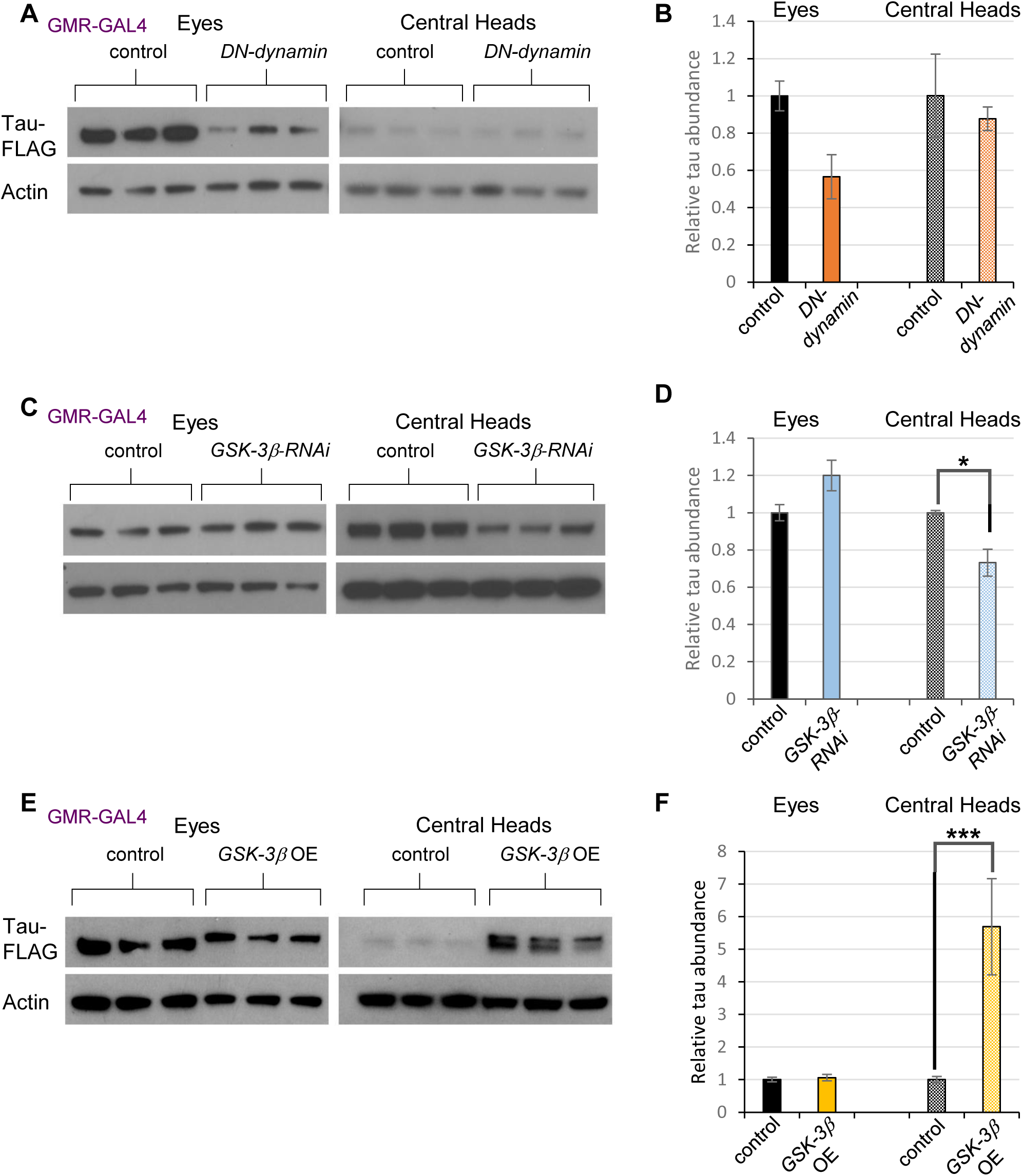
Dynamin influences tau spread in recipient tissues, while *GSK-3β* affects tau spread in source tissues. (A) Western blot analysis of protein extracts from the eyes and central heads of 20-day-old flies expressing both tau and dominant-negative (DN) dynamin in the eye. The experimental groups bore *GMR-QF2w* driving *tau-T2A-GFP* and *GMR-GAL4* driving a GAL4-responsive DN dynamin construct. Control flies lacked the transgene encoding DN dynamin. (B) Quantification of the data in panel A. (C) Western blot analysis of protein extracts from the eyes and central heads of 20-day-old flies expressing both tau and RNAi to *GSK-3β* in the eye. The experimental groups bore *GMR-QF2w* driving *tau-T2A-GFP* and *GMR-GAL4* driving a GAL4-responsive RNAi to *GSK-3β.* Control flies lacked the *GSK-3β* RNAi transgene. (D) Quantification of the data in panel C. (E) Western blot analysis of protein extracts from the eyes and central heads of 20-day-old flies expressing tau and overexpressing *GSK-3β* in the eye. The experimental groups bore *GMR-QF2w* driving *tau-T2A-GFP* and *GMR-GAL4* driving a GAL4-responsive *GSK-3β* construct. Control flies lacked the *GSK-3β* transgene. (F) Quantification of the data in panel E. For all experiments described in this figure, each condition was represented by three biological replicates. \**p* < 0.05, \*\*\**p* < 0.005 by Student *t-*test.

### Exploring the specificity of the tau spread modifiers

Many neurodegenerative disorders are characterized by brain protein aggregates that appear to spread between brain regions. This raises an important question: Are the mechanisms of toxic protein spread shared between neurodegenerative diseases, or are they disease specific? To begin to address this matter, we created a transgenic line that expresses α-synuclein protein, a major component of the Lewy body brain protein aggregates that are observed in Parkinson’s disease and related disorders (32, 41). Substantial evidence indicates that excess α-synuclein is toxic and that it has the ability to spread between brain regions (21, 41, 42). Our α-synuclein transgenic line was created in the same fashion as our tau transgenic line. Specifically, we created a Q-responsive transgenic construct that contained the coding sequences of α- synuclein and GFP with an intervening *T2A* cleavage peptide (Fig. 7A). Driving this transgene in the eye with *GMR-QF2w* produced the expected expression of α-synuclein and GFP (Fig. 7B). Alpha-synuclein was also detected in central heads, but GFP was not, indicating that α- synuclein spreads beyond its site of expression as tau does (Fig. 7B). However, unlike tau, the abundance of α-synuclein in central heads did not increase with adult age (Fig. 7C), perhaps indicating maximum spread of α-synuclein occurs during the pupal stage of development.

**Figure 7.**
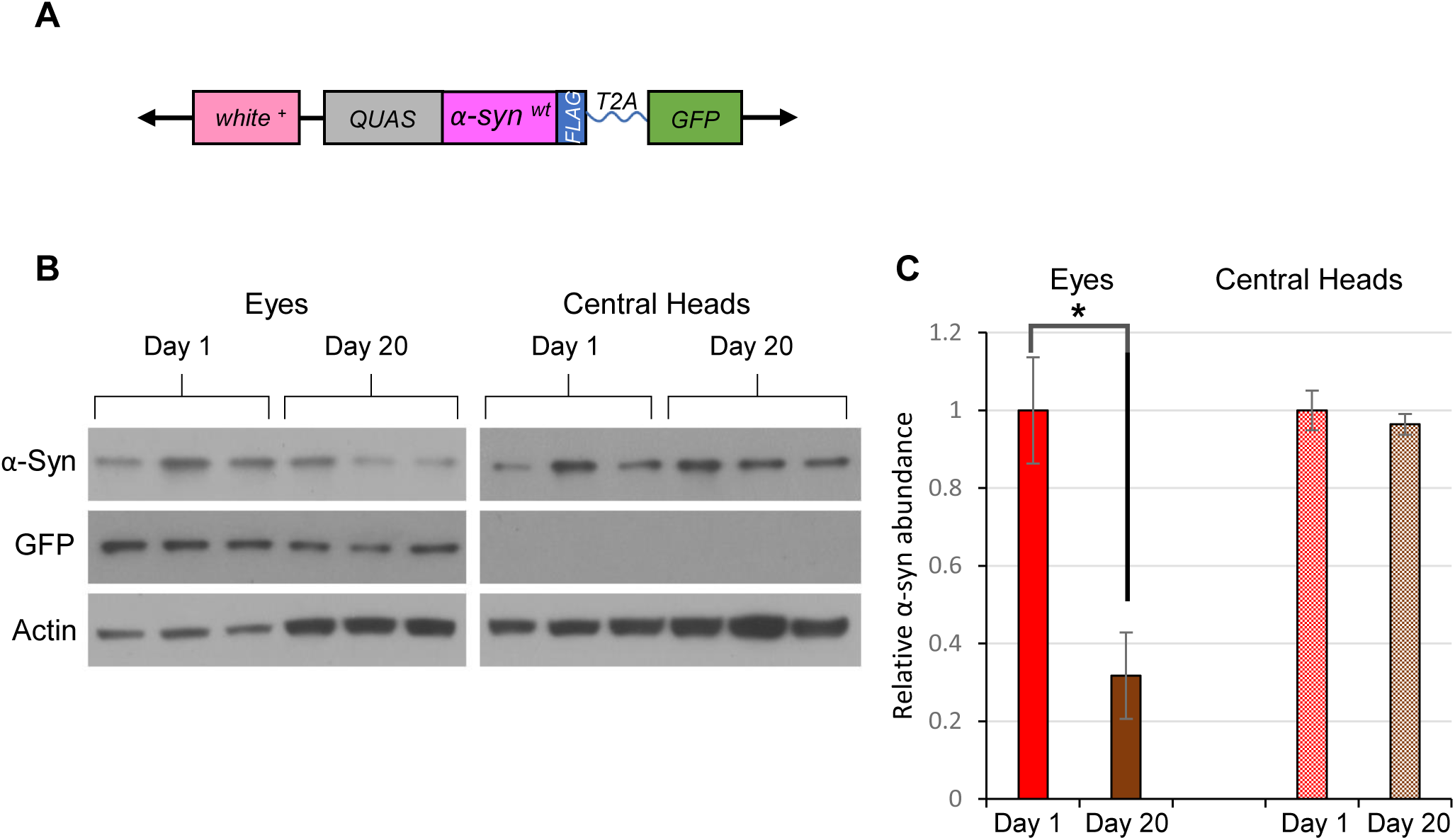
Alpha-synuclein expressed in the fly eye spreads to the brain, but spread is not progressive with age. (A) Schematic description of the α*-synuclein-T2A-GFP* transgene (see Fig. 1A). (B) Western blot analysis using antisera against FLAG, GFP, and actin, performed on protein extracts from the eyes and central heads of 1-day-old and 20-day-old flies expressing α- synuclein in the eye. Flies bore the *GMR-QF2w* driver and the *α-synuclein-T2A-GFP* transgene. Each condition was represented by three biological replicates. (C) Quantification of α-synuclein abundance using the data in panel B. \**p* < 0.05 by Student t test.

We then tested whether factors that influenced tau spread also influenced α-synuclein spread. Specifically, we repeated our experiments with the RNAi targeting *GSK-3β* and the dominant-negative dynamin construct in flies expressing our *α-synuclein-T2A-GFP* construct. These perturbations had no significant effect on the spread of α-synuclein (Fig. 8A-D). When we overexpressed GSK-3β, we found an increase in α-synuclein spread (Fig 8E, F), but the effect of this manipulation on α-synuclein was considerably smaller than its previous effect on tau (Fig. 3). Overall, our findings indicate that most of the genetic perturbations that influence tau spread were specific to tau.

**Figure 8.**
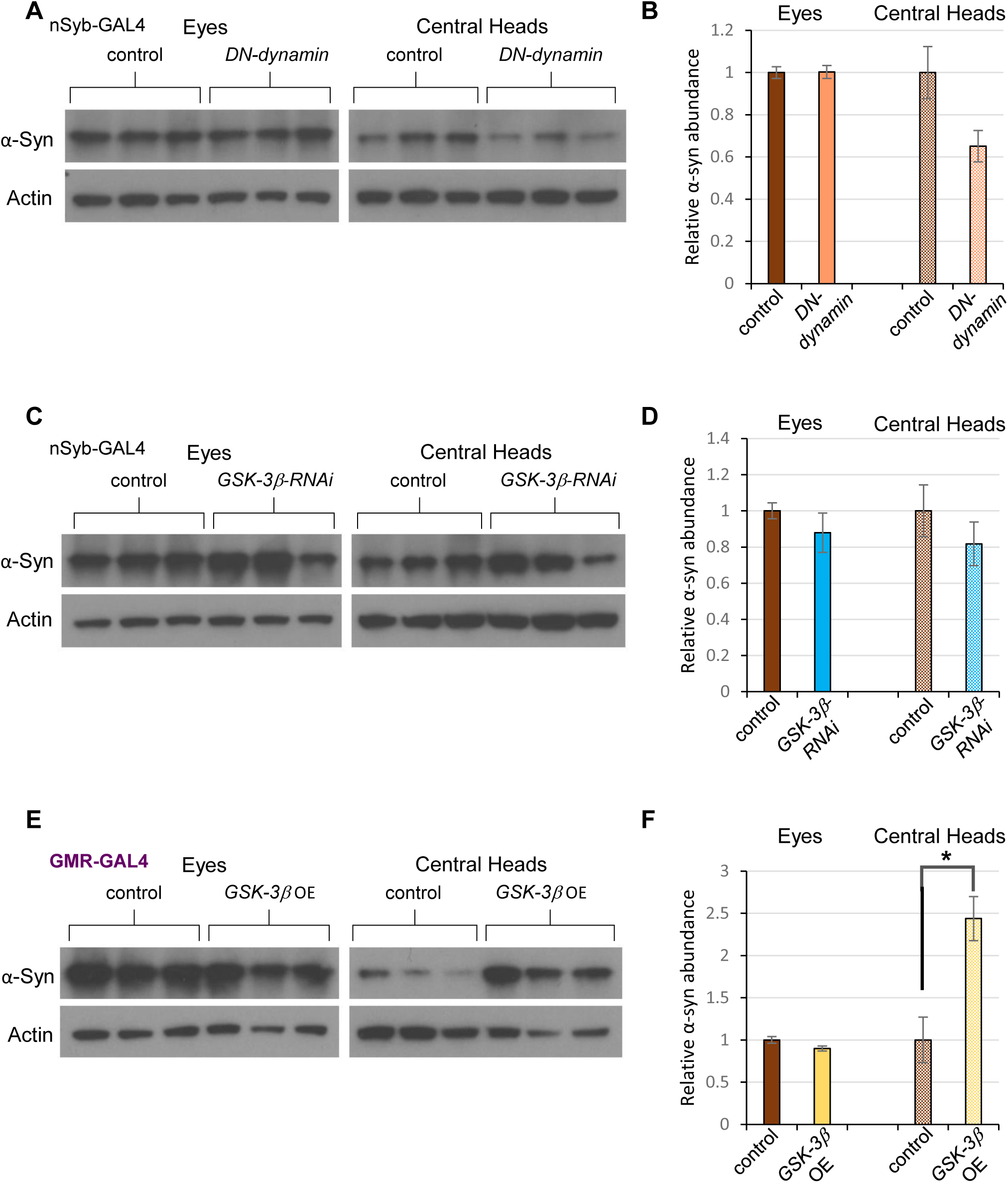
tau spread modifiers have little or no influence on α-synuclein spread. (A) Western blot analysis of protein extracts from the eyes and central heads of 20-day-old flies expressing α-synuclein in the eye and a dominant-negative (DN) form of dynamin in neurons. The experimental group bore *GMR-QF2w* driving *α-synuclein-T2A-GFP* and *nSyb-GAL4* driving a dominant-negative dynamin. Control flies lacked the transgene encoding the DN form of dynamin. (B) Quantification of the data in panel A. (C) Western blot analysis of protein extracts from the eyes and central heads of 20-day-old flies expressing α-synuclein in the eye and RNAi to *GSK-3β* in neurons. Specifically, *GMR-QF2w* drove *α-synuclein-T2A-GFP*, and the pan-neuronal driver *nSyb-GAL4* drove a GAL4-responsive RNAi targeting *GSK-3β*. Control flies lacked the *GSK-3β* RNAi transgene. (D) Quantification of the data in panel C. (E) Western blot analysis of protein extracts from the eyes and central heads of 20-day-old flies expressing *α- syn-T2A-GFP* in the eye using *GMR-QF2w* and co-expressing a GAL4-responsive *GSK-3β* transgene using *GMR-GAL4*. Control flies lacked the *GSK-3β* transgene. (F) Quantification of the data in panel E. For all experiments described in this figure, each condition was represented by three biological replicates. \**p* < 0.05 by Student *t*-test.

## DISCUSSION

The prion-like spread of toxic proteins from one brain region to another has gained increasing acceptance as an important pathological feature of many neurodegenerative diseases (43–46). Over the past several years, this process has been recapitulated in a number of model systems, including *Drosophila* (26, 28, 29). However, this recent work has several limitations. First, it can be challenging to distinguish between spread of the toxic protein and mild misexpression of the gene encoding the toxic protein outside of the designated target tissues. Second, vertebrate systems are not suitable for rapid exploration of candidate pathways because of the time involved in generating and aging the animals required. Third, tissue dissection and confocal microscopy to assess the extent of toxic protein spread is both time consuming and not readily amenable to quantification, particularly in instances where the differences between genotypes are modest. Our approach overcomes all of these limitations by creating transgenic *Drosophila* expressing tau and GFP from the same transgene and using a simple western blot procedure to detect the migration of tau. Moreover, because our system makes use of the Q expression system to drive tau expression, the many existing GAL4-responsive reagents can be used to inactivate and overexpress candidate spread modifiers in virtually any desired tissue without influencing tau expression.

Our work is supported by previous evidence that *Drosophila* can be used to study the mechanisms underlying the spread of neurodegeneration-associated proteins (26, 28, 29). We now provide a set of tools for systematic study of this phenomenon. Importantly, several of our observations indicate that the spread of tau in *Drosophila* reflects the processes involved in the spread of tau in the human brain. First, the spread of tau does not appear to be a secondary consequence of loss of cellular integrity due to tau toxicity; if this were the case, GFP would also spread. Second, the fact that GFP does not spread is not due to the smaller size of GFP compared to tau. Our *α-synuclein-T2A-GFP* flies also show spread of their disease-associated protein, and α-synuclein is not larger but smaller than GFP. Third, age-dependent differences in the spread of tau and α-synuclein oppose the idea that spread could be a common artifact of our model system, and imply that tau and α-synuclein spread through distinct mechanisms. Further support for the idea of distinct spread mechanisms comes from the finding that most of the genetic perturbations that affected tau spread in our experiments did not detectably affect α- synuclein spread. Finally, our findings on specific candidate spread markers are consistent with work in other model systems, including vertebrates.

Our work also points to potential mechanisms underlying the spread of tau protein. Specifically, we found that genetic perturbations that reduce or eliminate the activity of *Drosophila* dynamin or GSK-3β resulted in reduced tau spread. Both findings are consistent with previous work in *Drosophila* and mammals (26, 28, 47, 48). For example, a *Drosophila* model of huntingtin protein toxicity demonstrated that selectively blocking endocytosis, using the same dominant-negative dynamin construct used in our work, prevented non–cell autonomous huntingtin-mediated degeneration (28). The influence of dynamin on huntingtin protein toxicity was seen in the tissues that acquired huntingtin protein from surrounding tissues (28), consistent with our observations showing that pan-neuronal inactivation of dynamin reduced tau spread but selective inhibition of dynamin in the eye did not. Of even greater relevance to our current work, a recent *Drosophila* study showed that *GSK-3β* inactivation reduced tau spread from the eye to the brain (26). However, while the previous work used the GAL4 system to express both tau and the RNAi targeting *GSK-3β*, our system allowed us to manipulate GSK-3β activity independently of tau. We were therefore able to show that GSK-3β is selectively required in the donor tissue to influence tau spread. Furthermore, although we did not directly test phosphorylation levels, our findings suggested that the effect of overexpressing GSK-3β was at least partly mediated by hyperphosphorylation of tau. Previous work has shown that tau is a direct substrate of GSK-3β (36) and that phosphorylation increases tau toxicity (35). Our experiments showed that overexpression of GSK-3β caused a mobility shift for tau protein that was consistent with increased phosphorylation. Together, these findings strongly imply that the influence of GSK-3β on tau spread is at least partially a consequence of tau hyperphosphorylation. The mechanism by which tau hyperphosphorylation would influence its spread is unclear from our work, but one previously proposed explanation is that the reduced affinity of phosphorylated tau for microtubules might trigger its misfolding and secretion through a non-canonical secretory mechanism (49).

Previous work has also shown that GSK-3β can phosphorylate α-synuclein on Serine 129 (50) and that Serine 129 phosphorylation increases α-synuclein toxicity in multiple model systems (51–53). Our finding that GSK-3β overexpression increases α-synuclein spread raises the possibility that α-synuclein is more toxic upon phosphorylation of Serine 129 because this phosphorylation event alters α-synuclein spread. However, as GSK-3β overexpression had no obvious influence on α-synuclein mobility in our western blot assays, our findings are also consistent with the possibility that the influence of GSK-3β on α-synuclein spread is indirect. Although GSK-3β overexpression does affect tau mobility, indicating that tau is likely a direct target GSK-3β activity, the influence of GSK-3β on tau spread may also be at least partially indirect. Our new model system will facilitate efforts to resolve these and other questions.

A strength of our approach is its ability to detect and rule out candidate spread modifiers that alter overall tau abundance. For instance, we found that NSF knockdown caused a reduction of tau abundance in the brain, consistent with an earlier report that the same manipulation reduced huntingtin spread in *Drosophila* (28). If not for the simplicity and sensitivity of detecting differences in tau abundance within the source tissue (eye) afforded by our methodology, we would have classified NSF as a modifier of tau spread rather than a modifier of tau abundance. Furthermore, our work on NSF revealed another useful feature of our system. We found that the effect of NSF knockdown on tau abundance was specific to tau, because GFP expressed from the same transgene was not reduced in abundance by this manipulation (Fig. S1). Together these findings indicate that the effect of NSF knockdown on tau abundance is not an artifact of the expression systems we are using, or a general defect in expression, but rather a specific effect of NSF on tau abundance. While we cannot at present explain how reduced NSF activity selectively alters tau abundance, previous work has shown that NSF interacts physically (54) and genetically (55) with cytoskeletal components. Furthermore, tau itself has recently been shown to interact with NSF and reduce its activity (56). Regardless of how NSF alters tau abundance, these results further demonstrate the utility and power of our model to identify and categorize tau modifiers.

Over the past 20 years, researchers have identified hundreds of genetic loci that influence the risk of AD, PD, and other neurodegenerative disorders associated with the accumulation and spread of protein aggregates in the brain (57–64). However, for many of these loci we do not know how they influence disease risk, or precisely which gene in the linkage region is the risk-modifying factor. While these loci likely influence risk in a variety of ways, we anticipate that at least some act by enhancing the spread of tau or α-synuclein to other brain regions. Thus, we anticipate that our model systems for identifying modifiers of tau and α-synuclein spread will be extremely useful in identifying genes that influence the risk of AD and PD, and the mechanisms by which they act. Knowledge acquired from these studies could ultimately lead to the development of therapeutic strategies designed to block the spread of tau and α-synuclein, and thus to prevent or slow development of neurodegenerative disease.

## MATERIALS AND METHODS

### Fly stocks and generation of transgenic lines

All fly stocks were maintained on regular cornmeal-molasses food at 25°C using a 12-h light/dark cycle. The *GMR-QF2w* (BDSC 59283), *nSyb-GAL4* (51941), and *GMR-GAL4* (8121) driver stocks were obtained from the Bloomington Drosophila Stock Center. Stocks bearing RNAi constructs targeting potential spread modifiers were also acquired from BDSC: *UAS-sgg-RNAi* (31309) and *UAS-sgg* (5361), identified in text by the name of the mammalian ortholog *GSK3β*; *UAS-shi^ts^* (44222; *dynamin*); and *UAS-comatose-RNAi* (31666; *NSF*).

To produce the transgenic fly lines created for this study, we first generated two recombinant constructs in the pQUAS_WALIUM20 plasmid, one with wild-type (2N4R) human tau sequence and the other with wild-type α-synuclein sequence (see Fig. 1A). Both disease-associated proteins were thus placed under QUAS promoter control. The sequence of tau or α-synuclein was followed by a FLAG sequence, a cleavable *T2A* domain, and then the coding sequence of GFP. These constructs were then injected into *Drosophila* embryos to generate transgenic flies with the assistance of Rainbow Transgenic Flies, Inc.

### Immunohistochemistry

Brains from 40-day old female flies were dissected in cold 1X phosphate-buffered saline (PBS, pH 7.5), and fixed in 4% paraformaldehyde/PBS for 45 min. Tissues were washed in 1x PBS containing 0.1% Triton X-100. Fixed brains were probed with 1:500 rabbit anti-FLAG and 1:800 mouse anti-GFP primary antibodies overnight at 4°C, followed by staining with anti-mouse Alexa 488 and anti-rabbit Alexa 568 antibodies. Samples were mounted using ProLong Gold Antifade (Molecular Probes, P10144), and imaging was done using an SP8 Confocal Microscope (Leica).

### Dissections and western blotting

Groups of 18 to 20 female flies were harvested by flash freezing at appropriate time points, most at day 20-21, with age matched controls. Age is indicated in the legend of each figure. Under a dissecting microscope, eyes were separated from central heads using a razor blade. Sets of eyes and sets of central heads were homogenized separately, each in 100 µL of 1X RIPA lysis buffer (50 mM Tris, pH 7.4; 150 mM NaCl; 1% Nonidet P-40; 0.5% sodium deoxycholate; 0.1% SDS). Proteins were separated by SDS-PAGE using 4%-20% MOPS-acrylamide gels (GenScript Express Plus M42012) and transferred electrophoretically onto Immobilon PVDF membrane (Merck). The membrane was transferred to blocking buffer (5% nonfat dry milk in 1x PBS with 0.1% Tween-20) for 1 h. Membranes were incubated overnight with primary antibodies diluted in blocking buffer. Primary antibody dilutions were as follows: 1:1000 rabbit anti-FLAG (Cell Signaling Technologies 14793S), 1:1000 anti-alpha-synuclein (BD Transduction Laboratories 610787), 1:500 mouse anti-GFP (Biolegend 668205) and 1:5000 mouse anti–β-actin (Millipore Sigma MAB1501). After three washes, the membrane was incubated with secondary antibody diluted 1:7500 in blocking buffer (anti-rabbit, anti-mouse HRP, Bio-Rad). Signal was detected using Pierce ECL Western Blotting Substrate (Fisher 32106). Densitometric quantitation of western blots was performed using Fiji software (NIH) by an investigator blinded to genotype. Signal from the protein of interest was normalized to actin levels.

### Statistical analysis

All experiments were performed at least three times (n ≥ 3). Densitometric values were normalized to actin and log-transformed to stabilize the variance before means were compared using Student *t*-test.

## Acknowledgments

The authors gratefully acknowledge the assistance of Dirk Hueglin and Eurofins Genomics Blue Heron LLC in the generation of transgenic constructs, and the assistance of Rainbow Transgenic Flies, Inc., with embryo injection.

## Author contributions

Conceptualization: L.J.P., R.E.T., K.B., E.S.V.; Methodology: L.J.P., R.E.T., E.S.V.; Validation: R.E.T., K.B., G.M., L.V.F.; Formal analysis: R.E.T., K.B.; Investigation: R.E.T., K.B., G.M., L.V.F.; Data curation: R.E.T., K.B., G.M.; L.V.F.; Writing (original draft): K.B., L.J.P., R.E.T.; Writing (review & editing): E.S.V., R.E.T., K.B., L.J.P.; Visualization: K.B., R.E.T., E.S.V.; Supervision: L.J.P.; Project administration: R.E.T., L.J.P.; Funding acquisition: L.J.P, R.E.T., E.S.V.

## Data availability

All relevant data can be found within the article and its supplementary information.

---

**Supplemental Figure 1. GFP abundance is unchanged upon inactivation of NSF.** (A) Western blot analysis of protein extracts from the eyes of 30-day-old flies expressing *tau-T2A-GFP* in the eye and an RNAi targeting *NSF* in neurons using anti-GFP. The experimental group bore *GMR-QF2w* driving *tau-T2A-GFP* and *nSyb-GAL4* driving a GAL4-responsive RNAi to *NSF*. Control flies lacked the RNAi transgene. Each condition was represented by three biological replicates. (B) Quantification of the data in panel A.

